# Noise decorrelation optimizes SNR of GABA-edited MRS data: A comparison of RF coil combination methods

**DOI:** 10.1101/2025.03.05.637500

**Authors:** Amy E. Bouchard, Mark Mikkelsen

**Affiliations:** Department of Radiology, Weill Cornell Medicine, New York, NY, United States

**Keywords:** Coil combination, GABA, generalized least squares, magnetic resonance spectroscopy, phased-array coil, signal-to-noise ratio

## Abstract

**Introduction:** Determining the best radiofrequency (RF) coil combination method is crucial for maximizing the signal-to-noise ratio (SNR) to detect low concentration metabolites (e.g., γ-aminobutyric acid (GABA)) in magnetic resonance spectroscopy (MRS). We hypothesized that algorithms accounting for noise correlations between coil elements would optimize SNR, given that phased-array coils provide better SNR than surface coils and allow accelerated acquisitions, and methods accounting for noise correlations outperform those assuming no correlations.

**Methods:** We examined six coil combination methods, the latter half accounting for noise correlations: 1) equal weighting; 2) signal weighting; 3) S/N^2^ weighting; 4) noise-decorrelated combination (nd-comb); 5) whitened singular value decomposition (WSVD); 6) generalized least squares (GLS). We utilized MEGA-PRESS data from 119 participants (mean age: 26.4 ± 1 SD 4.2 years; males/females: 54/65) acquired on 3T GE and Siemens MRI scanners at 11 research sites, obtained from the Big GABA study. We measured the SNR of GABA and *N*-acetylaspartate (NAA). We also calculated the intersubject coefficients of variation of GABA.

**Results:** There were significant differences in SNR between coil combination methods for both GABA+ and NAA. More specifically, the noise decorrelation methods produced higher GABA+ and NAA SNR than the other approaches, where nd-comb, WSVD, and GLS produced, on average, ∼37% and ∼34% more SNR than equal weighting, respectively. GLS produced the highest SNR for GABA+ and NAA. The coefficients of variation for GABA+ were generally slightly smaller for the noise decorrelation methods.

**Conclusion:** Noise-decorrelation methods produced higher SNR than other methods, especially GLS, which should be investigated in advanced editing protocols.

## 1. Introduction

Magnetic resonance spectroscopy (MRS) is a noninvasive method to detect metabolites of interest in vivo. In MRS, there is an important need to obtain as much signal-to-noise ratio (SNR) as possible, given the considerably lower signal strengths of metabolites in the human body and brain. Improving SNR is critical for data reliability, quantification, and avoiding overly long scan durations, particularly for molecules in the ∼1–3 mM range such as γ-aminobutyric acid (GABA). Doing so would help MRS to become one step closer to potentially being used as a clinical tool. Combining signals using phased-array radiofrequency (RF) coils is effective for improving SNR in MRS (e.g., [1]). In addition, phased-array RF coils provide higher SNR than surface coils. They also allow for accelerated acquisitions through techniques such as GeneRalized Autocalibrating Partial Parallel Acquisition (GRAPPA) and SENSitivity Encoding (SENSE). However, it remains to be seen which coil combination methods lead to higher SNR than others.

There are multiple methods for combining phased-array RF coils to improve SNR, such as equal weighting, signal weighting, and S/N^2^ weighting. Equal weighting consists of equally summing over all channels after applying phasing based on the first point of a reference free induction decay (FID). Signal weighting corresponds to weighting based on the magnitude of a reference signal [2]. In addition, S/N^2^ weighting involves weighting centered on the ratio of signal to the square of the noise of a reference signal [3].

Considering that noise correlations between coils have been found to occur [4, 5], several noise decorrelation methods have also been developed, notably, noise decorrelated combination, whitened singular decomposition, and generalized least squares. Noise-decorrelated combination (nd-comb) consists of combining using principal component analysis (PCA)-based noise decorrelation and weighting by the signal-to-noise ratio of a reference signal [6]. Also, whitened singular value decomposition (WSVD) involves combining using PCA-based noise decorrelation and deriving the optimal signal through singular value decomposition of the noise-whitened FIDs [7]. Lastly, generalized least squares (GLS) implicates combining that is centered on generalized least squares where some degree of correlation in the noise is assumed [8].

Previous comparisons between coil combination methods have demonstrated that those that account for noise correlations between coil elements outperform methods that assume uncorrelated noise in brain [9], heart [10], and breast cancer tissue [11]. To the best of our knowledge, only one study investigated noise decorrelation methods in the brain. This study found that adaptively optimized combination (AOC) (a noise decorrelated method similar to GLS, under the condition that the unsuppressed water peak is used as the reference [12]) led to higher SNR than methods that did not account for noise (i.e., equal weighting, signal weighting, S/N, S/N^2^) (although this depended on the location of the MRS voxel of interest) [9].

However, it is unknown whether accounting for noise correlations between coil elements leads to better SNR than methods that do not for MRS acquisitions detecting low-concentration metabolites in the brain (e.g., GABA) as well as high-concentration metabolites such as n-acetylaspartate (NAA). Hence, this study investigated the optimal coil combination method(s) for GABA-edited MEGA-PRESS data. Based on the current literature [9–11], we hypothesized that noise decorrelation methods would produce superior SNR for GABA and NAA.

## 2. Methods

### 2.1 Theory

The complex phased-array time-domain MRS signal *S_k_*(*t*) from the kth coil element can be formulated as:

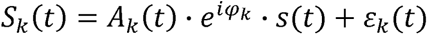

where *A_k_*(*t*) is the coil signal amplitude, ϕ*_k_* is the coil signal phase, *s*(*t*) is the FID signal, and ε*_k_*(*t*) is the coil noise. It is typical during coil combination to assume the noise between channels is uncorrelated. However, as previously stated, noise correlations are known to occur in multi-coil RF receive systems [4, 5] and should be accounted for during coil combination. To optimally combine the signals from each element, each *k*th signal must be appropriately weighted so that the combined signal is as follows:

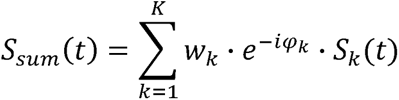

The goal is to find optimal weight factors *w_k_*. To determine the best approach, we assessed the following six coil combination methods: 1) equal weighting; 2) signal weighting [2]; 3) S/N^2^ weighting [3]; 4) nd-comb [6]; 5) WSVD [7]; and 6) GLS [8].

### 2.2 Participants

119 healthy adults (mean age: 26.4 ± 1 standard deviation (SD) (4.2 years)); male/female: 54/65), took part in this study. We obtained GABA-edited MEGA-PRESS data of these participants from the Big GABA study [13, 14] (https://www.nitrc.org/projects/biggaba/). Participants were free of neurological and psychiatric disorders. The Weill Cornell Medicine Institutional Review Board provided ethical approval for this study.

### 2.3 Data processing and analysis

We acquired data that were collected on 3T GE and Siemens scanners at 11 research sites [13, 14]. Single-voxel MEGA-PRESS data were acquired with the following parameters: TE/TR = 68/2000 ms; 320 transients; voxel size = 30 × 30 × 30 mm^3^. The MRS voxel was placed in the medial parietal lobe.

We estimated noise covariance matrices from the last quarter of either water-suppressed or -unsuppressed FIDs. We used 3.0 ppm Cr as a concentration reference. We processed data in Gannet (v. 3.2.1) [15] running on Matlab (v. R2024a). We calculated the SNR of the 3.0 ppm GABA+ peak in the difference spectrum and the 2.0 ppm NAA peak in the edit-OFF spectrum. We defined SNR as the ratio of the respective modeled signal amplitude divided by twice the SD of noise signal between 8 and 10 ppm in either the difference or OFF spectrum, respectively. We also calculated the means ± 1 SD for both GABA+/Cr and NAA/Cr. In addition, we calculated the intersubject coefficient of variation (CV) of GABA+/Cr ratios for each of the 11 datasets over the six approaches. We carried out statistical analyses using R (v4.4.0) through one-way repeated-measures ANOVA and post hoc comparisons (where appropriate).

## 3. Results

The mean (± 1 SD) GABA SNR for equal weighting, signal weighting, S/N^2^, nd-comb, WSVD, and GLS were, respectively: 17.6 ± 3.0; 20.9 ± 4.4; 20.9 ± 4.6; 23.8 ± 5.5; 24.0 ± 5.5; and 24.4 ± 6.1. There was a significant difference in GABA SNR between methods (*F*(1.7, 200.9) = 142.2, *p* < 0.0001). Post hoc comparisons revealed several between-method differences (*p* < 0.001; see **Table 1**). In particular, the noise-decorrelation methods produced higher GABA SNR than the other approaches (see **Figures 1** and **2**), with nd-comb, WSVD, and GLS producing, on average, ∼37% more SNR than equal weighting (see **Figure 3**). In addition, the group variability in GABA+/Cr ratios did not differ markedly between equal weighting, signal weighting, and S/N^2^ weighting (see **Figure 4**). On the other hand, there was a difference between the noise-decorrelation methods, where GLS produced the smallest dispersion, followed by WSVD and nd-comb (see **Figure 4**). The CV values of GABA+/Cr for equal weighting, signal weighting, S/N^2^, nd-comb, WSVD, and GLS, were respectively: 12.5%; 11.4%; 11.1%; 10.6%; 11.1%; and 10.9%.

**Figure 1.**
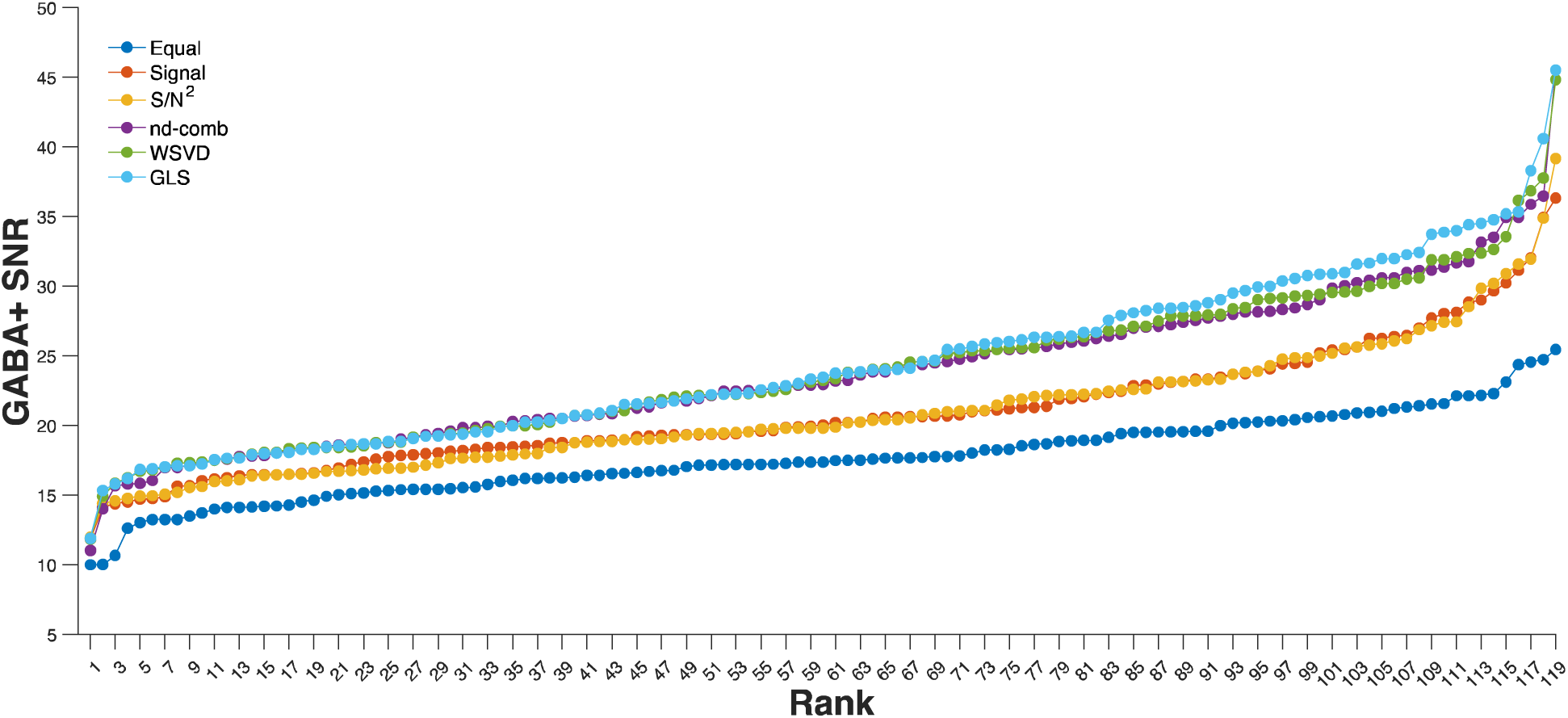
The SNR of GABA+ for all datasets using the six coil combination methods arranged in rank order.

**Figure 2.**
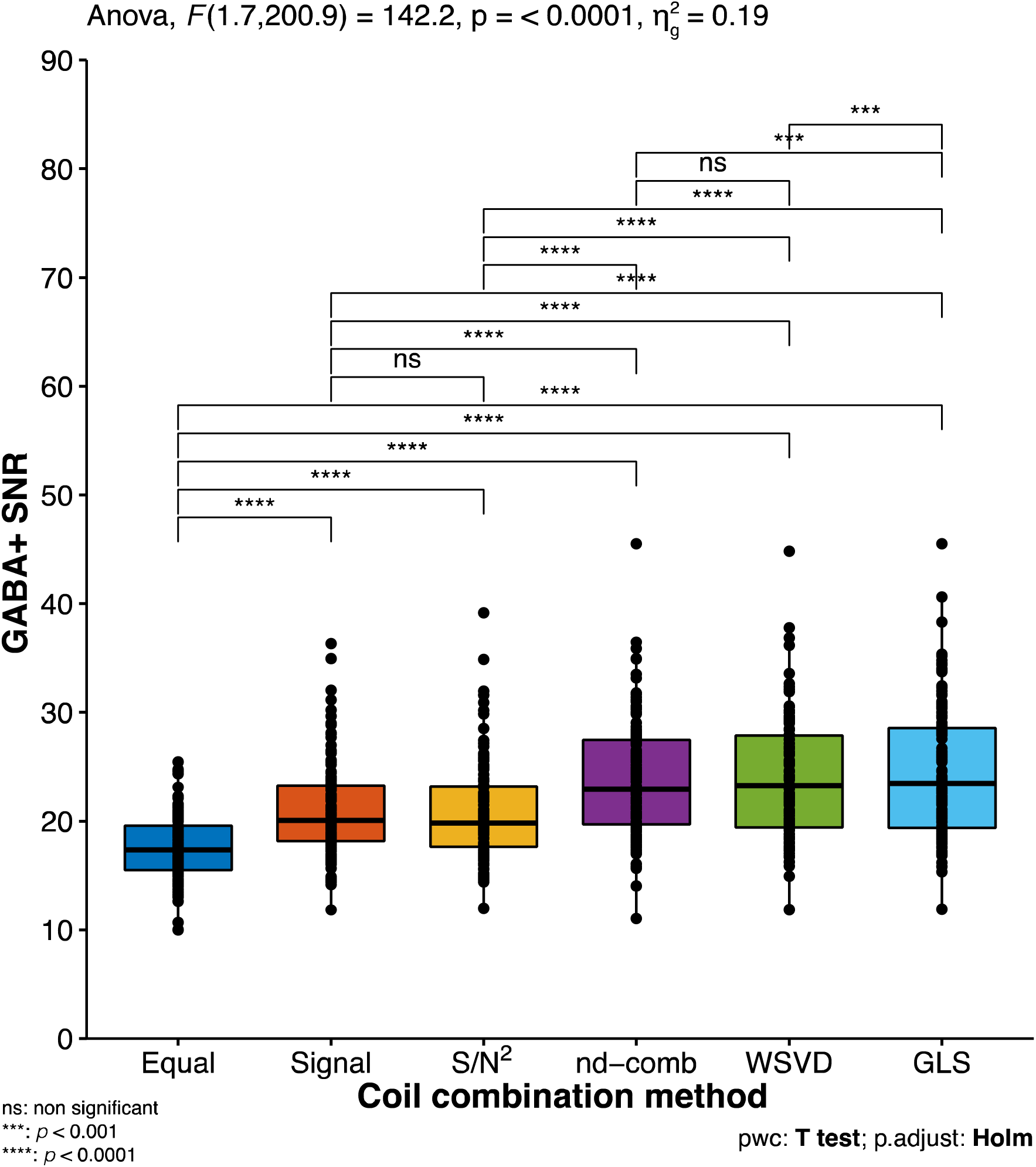
Boxplots for the six coil combination methods and post hoc group comparisons for GABA+.

**Figure 3.**
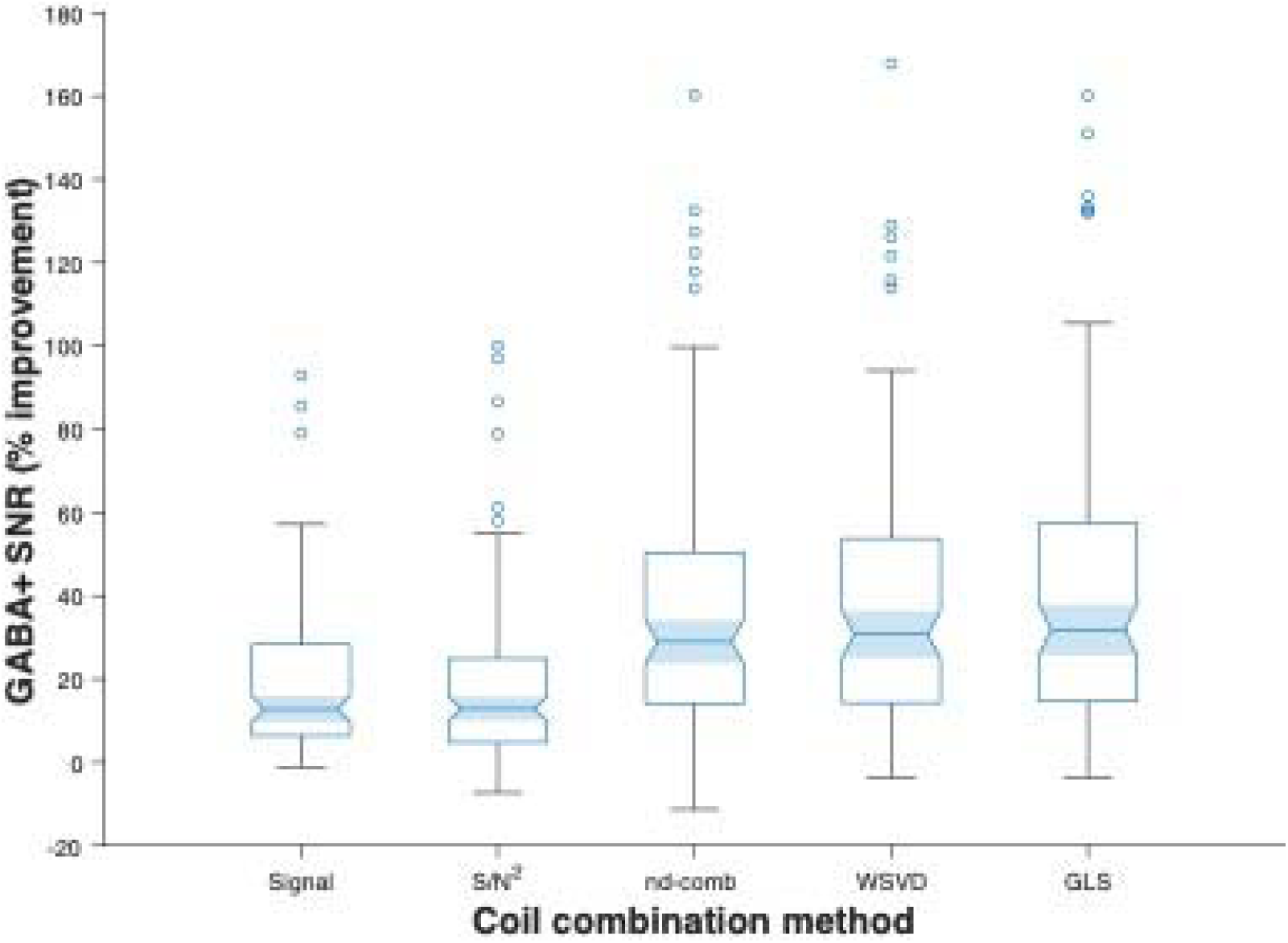
Boxplots of percentage improvement of each coil combination method in GABA+ SNR relative to equal coil combination weighting.

**Figure 4.**
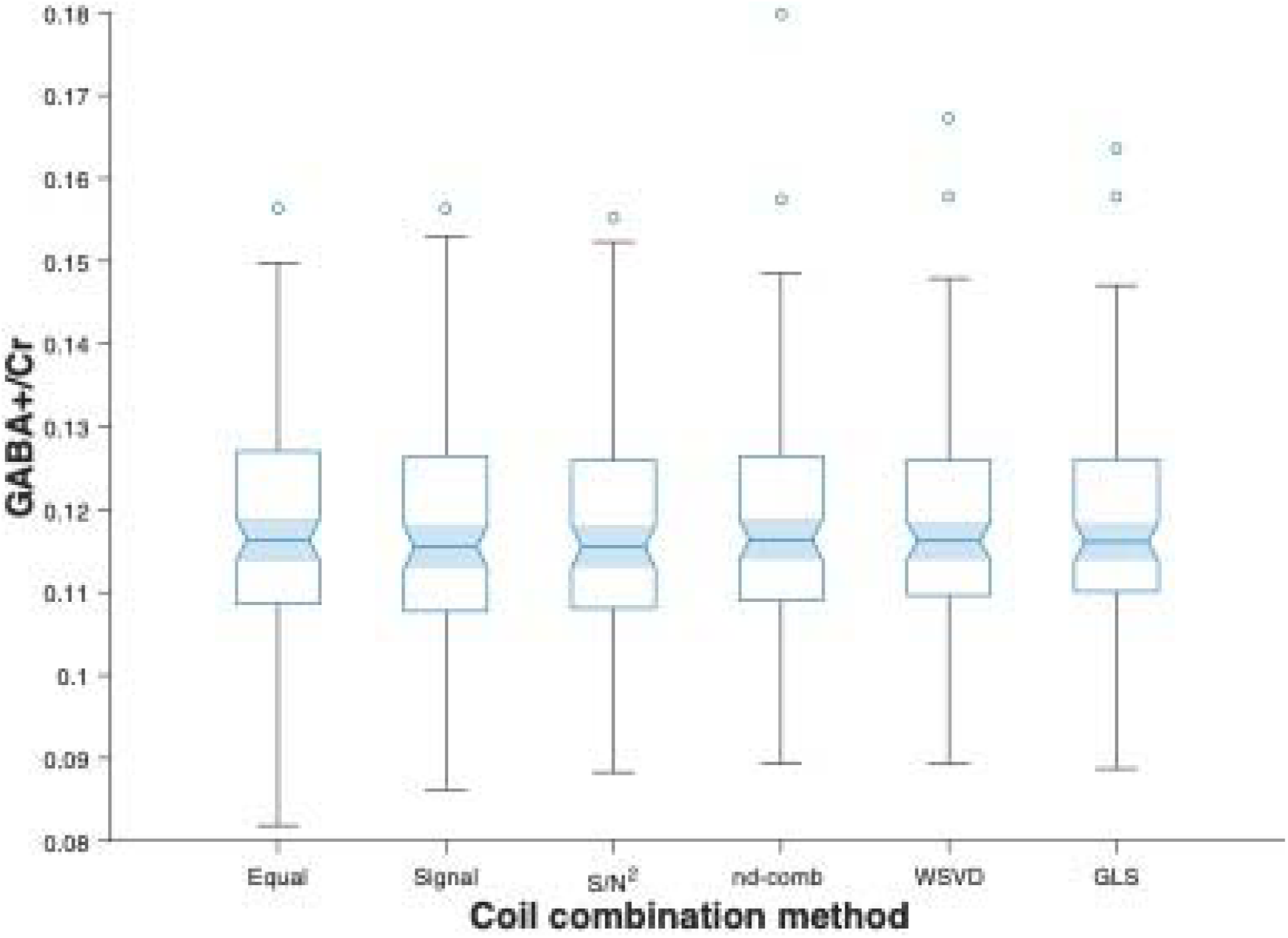
Boxplots of the quantitated GABA+/Cr ratios based on each coil combination method.

**Table 1.** Statistics (ANOVA and post hoc comparisons) for GABA+ and NAA SNR.

|  |  |  |  |  |  |
| --- | --- | --- | --- | --- | --- |
| GABA+ |  |  |  |  |  |
| ANOVA | Dfn | Dfd | *F* statistic |  | *p*-value |
|  | 1.7 | 200.9 | 142.2 |  | < 0.0001 |
| Post hoc pairwise comparisons |  |  |  |  |  |
| Group 1 | Group 2 | *t*-statistic | df | *p*-value | Adjusted *p*-value |
| Equal | Signal | -11.5 | 118 | < 0.0001 | < 0.0001 |
| Equal | S/N<sup>2</sup> | -10.2 | 118 | < 0.0001 | < 0.0001 |
| Equal | nd-comb | -13.7 | 118 | < 0.0001 | < 0.0001 |
| Equal | WSVD | -14.0 | 118 | < 0.0001 | < 0.0001 |
| Equal | GLS | -13.6 | 118 | < 0.0001 | < 0.0001 |
| Signal | S/N<sup>2</sup> | 0.5 | 118 | 0.6 | 0.6 |
| nd-comb | Signal | 9.6 | 118 | < 0.0001 | < 0.0001 |
| Signal | WSVD | -10.6 | 118 | < 0.0001 | < 0.0001 |
| GLS | Signal | 10.7 | 118 | < 0.0001 | < 0.0001 |
| nd-comb | S/N<sup>2</sup> | 10.0 | 118 | < 0.0001 | < 0.0001 |
| S/N<sup>2</sup> | WSVD | -11.2 | 118 | < 0.0001 | < 0.0001 |
| GLS | S/N<sup>2</sup> | 11.1 | 118 | < 0.0001 | < 0.0001 |
| nd-comb | WSVD | -1.0 | 118 | 0.3 | 0.6 |
| GLS | nd-comb | 3.9 | 118 | < 0.001 | < 0.001 |
| GLS | WSVD | 3.7 | 118 | < 0.001 | < 0.001 |
| NAA |  |  |  |  |  |
| ANOVA | Dfn | Dfd | *F* statistic |  | *p*-value |
|  | 1.5 | 171.9 | 97.0 |  | < 0.0001 |
| Post hoc pairwise comparisons |  |  |  |  |  |
| Group 1 | Group 2 | *t*-statistic | df | *p*-value | Adjusted *p*-value |
| Equal | Signal | -10.9 | 118 | < 0.0001 | < 0.0001 |
| Equal | S/N<sup>2</sup> | -9.4 | 118 | < 0.0001 | < 0.0001 |
| Equal | nd-comb | -10.7 | 118 | < 0.0001 | < 0.0001 |
| Equal | WSVD | -11.1 | 118 | < 0.0001 | < 0.0001 |
| Equal | GLS | -11.0 | 118 | < 0.0001 | < 0.0001 |
| Signal | S/N<sup>2</sup> | 0.9 | 118 | 0.3 | 0.7 |
| nd-comb | Signal | 7.5 | 118 | < 0.0001 | < 0.0001 |
| Signal | WSVD | -8.3 | 118 | < 0.0001 | < 0.0001 |
| GLS | Signal | 8.9 | 118 | < 0.0001 | < 0.0001 |
| nd-comb | S/N<sup>2</sup> | 8.4 | 118 | < 0.0001 | < 0.0001 |
| S/N<sup>2</sup> | WSVD | -8.8 | 118 | < 0.0001 | < 0.0001 |
| GLS | S/N<sup>2</sup> | 9.4 | 118 | < 0.0001 | < 0.0001 |
| nd-comb | WSVD | 0.7 | 118 | 0.5 | 0.7 |
| GLS | nd-comb | 4.2 | 118 | < 0.0001 | < 0.001 |
| GLS | WSVD | 5.1 | 118 | < 0.0001 | < 0.0001 |

ANOVA, analysis of variance; GLS, generalized least squares; nd-comb, noise-decorrelated combination; WSVD, whitened singular value decomposition

Furthermore, the mean (± 1 SD) NAA SNR for the six methods were, respectively: 305.7 ± 68.8; 363.6 ± 111.3; 361.0 ± 113.4; 407.7 ± 147.4; 405.9 ± 142.1; and 418.2 ± 156.2. There was a significant difference in NAA SNR between methods (*F*(1.5, 171.9) = 97.0, *p* < 0.0001). Post hoc analyses demonstrated numerous differences between groups (*p* < 0.001; see **Table 1**). More specifically, the methods that consider noise correlations produced higher NAA SNR than the other approaches (see **Figure 5** and **Figure 6**), with nd-comb, WSVD, and GLS yielding, on average, ∼34% more SNR than equal weighting (see **Figure 7**).

**Figure 5.**
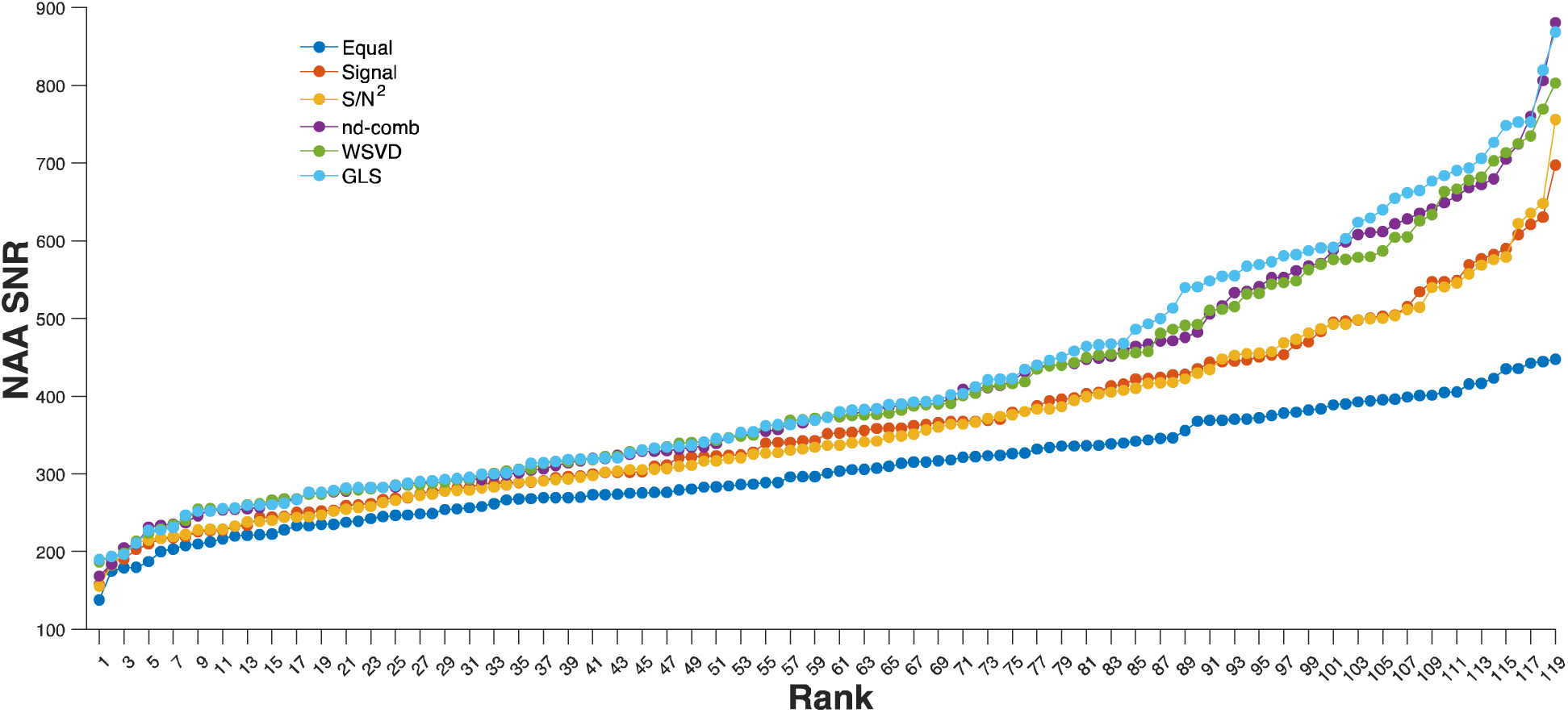
The NAA SNR of each participants’ data based on each of the six coil combination methods ranked by increasing SNR.

**Figure 6.**
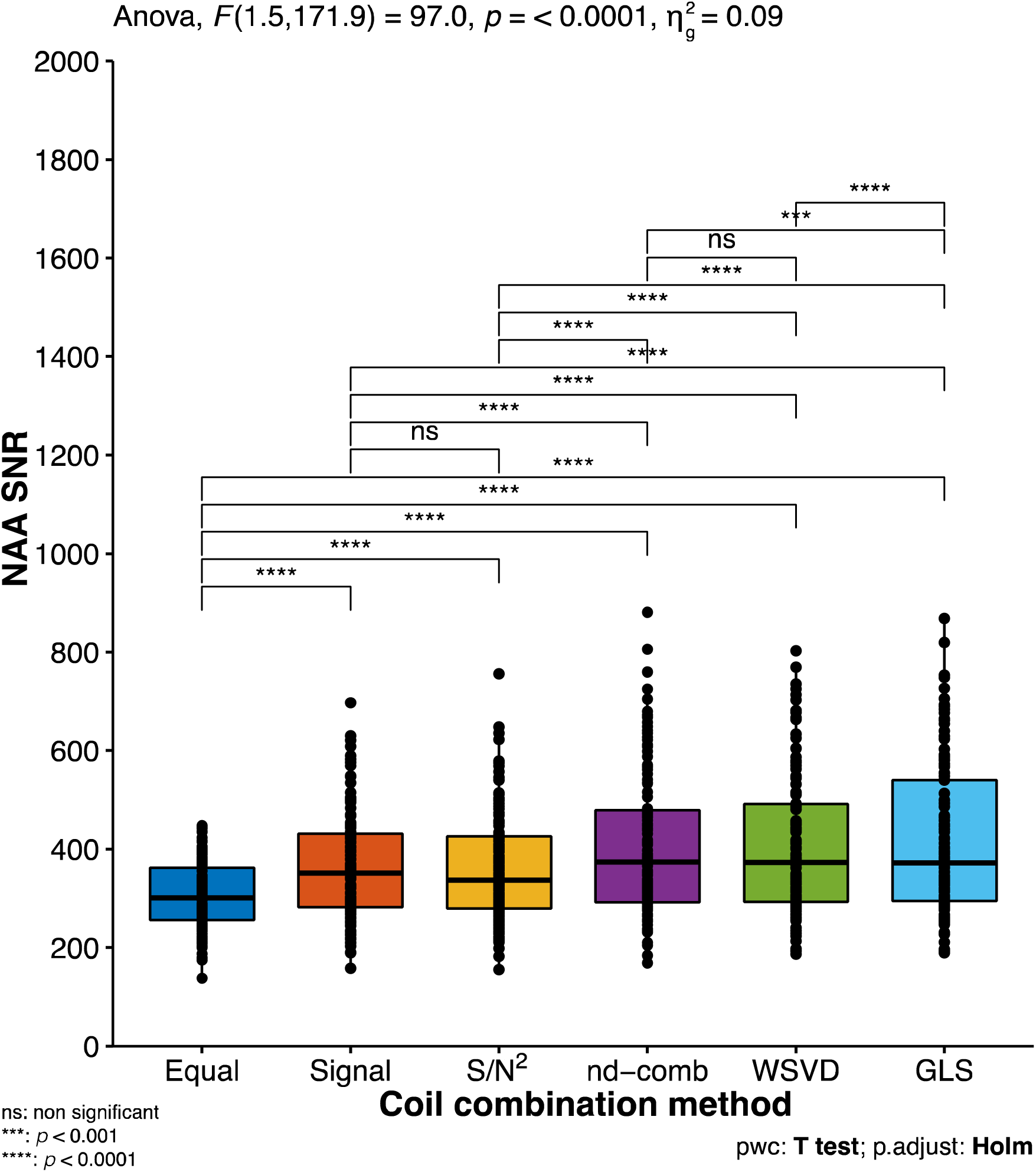
Boxplots for the six coil combination methods and post hoc group comparisons for NAA.

**Figure 7.**
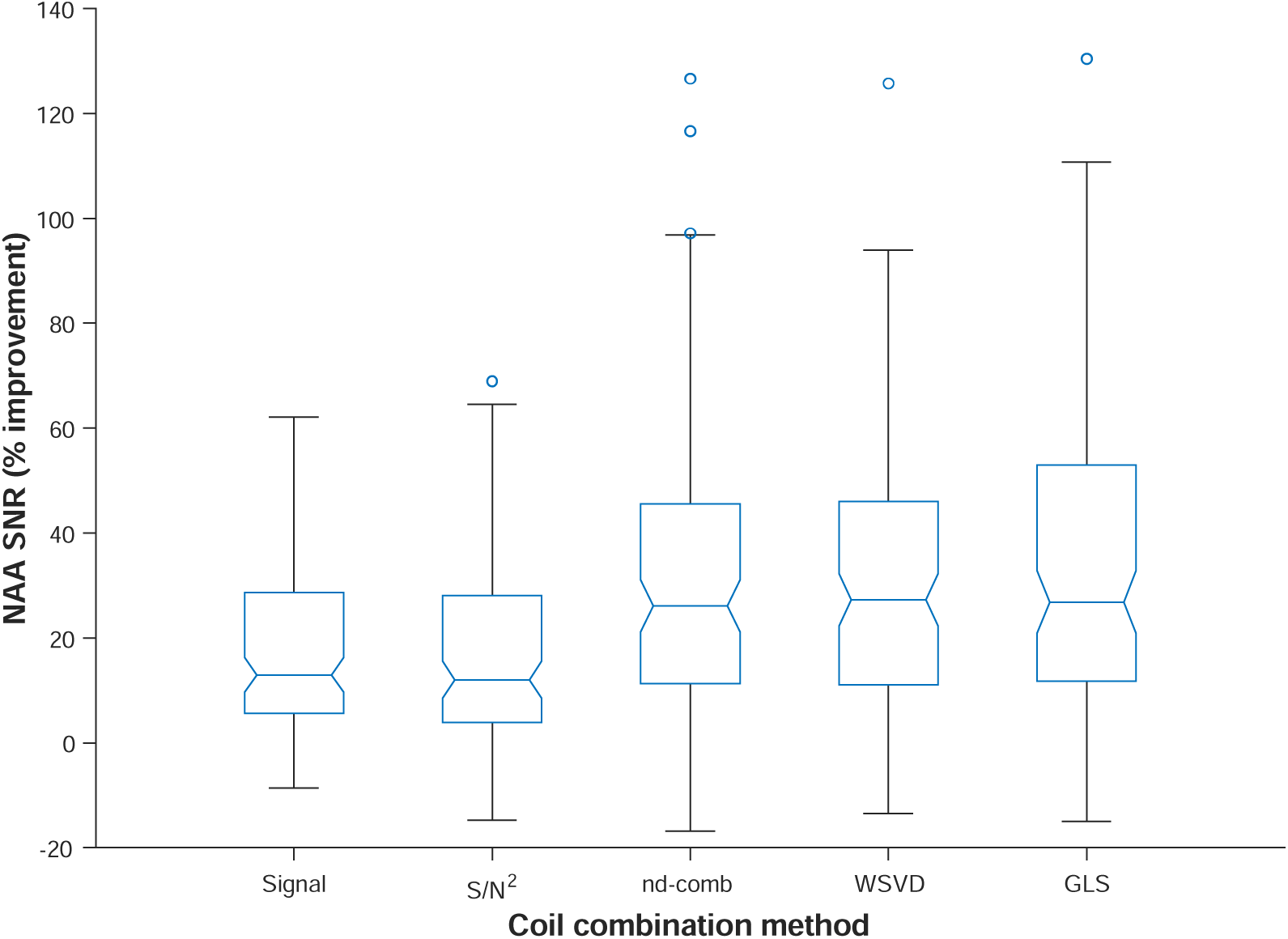
Boxplots of percentage improvement of each coil combination method in NAA SNR relative to equal coil combination weighting.

## 4. Discussion

This study aimed to evaluate whether and which RF coil combination method(s) improved SNR using GABA-edited MEGA-PRESS data. We hypothesized that noise decorrelation methods would show superior SNR for both GABA+ and NAA. As expected, our study showed that noise decorrelation methods, that is, nd-comb, WSVD, and GLS, produced superior SNR than the RF coil combination methods that do not account for noise correlations. This is in line with findings from the literature [9–11]. As previously outlined, Fang et al. (2016) discovered that AOC produced greater SNR than equal weighting, signal weighting, S/N, and S/N^2^ in the human brain [9]. Another study found that variations of WSVD combination methods led to higher SNR for human (cardiac spectra) experiments than methods that did not decorrelate noise (e.g., Roemer’s combination method) [10]. Likewise, one study found that for human breast tumor specimens, AOC resulted in superior SNR compared to other methods (e.g., equal weighting, nd-comb, signal weighting, S/N, and S/N^2^, although it performed similarly to WSVD) [11]. In addition, in patients with breast cancer, AOC led to the biggest SNR improvement, which was higher than all other methods except for nd-comb (AOC performed similarly to nd-comb).

Likewise, in healthy volunteers, AOC had the largest SNR improvement than all other methods except for WSVD (there was no difference between AOC and WSVD). Interestingly, in our study, there was no difference between WSVD and nd-comb for GABA and NAA SNR, yet GLS produced greater SNR than the two. Altogether, our findings showed that methods that accounted for noise decorrelation methods outperformed those that did not account for such correlations, which agrees with the literature.

Moreover, we found no clear advantage between noise decorrelation methods; that is, nd-comb, WSVD, and GLS all produced similar results, yet both GLS and nd-comb had the smallest CVs. This aligns with the study by Mallikourti et al. (2019), described above, in which AOC had high but similar improvements in SNR as nd-comb in patients with breast cancer. However, there was a slight advantage in our study for GLS in that it produced the highest SNR than the other noise decorrelation methods for both GABA+ and NAA, and one of the lowest CVs for GABA+ (nd-comb produced slightly smaller CV), as well as the smallest dispersion out of the noise decorrelation methods. There is mixed agreement within the literature. On the one hand, our finding opposes another study that found lower CVs for GLS than nd-comb in in vivo (human) experiments [8]; yet, on the other hand, in this same study [8], GLS produced lower CVs than WSVD, which agrees with our finding. It is possible that the differences between CVs are negligible and that simply using a noise-decorrelated method suffices, although finding the optimal noise-decorrelated coil combination strategy should be investigated in future work.

This study is not without limitations. Although we collected data from various scanners and two separate vendors, we are missing data for the Philips vendor, as the coil combinations from the Big GABA study were already combined. Future work examining coil combination methods collected on Phillips scanners should be conducted. In addition, we acknowledge that other noise decorrelations exist (e.g., [9, 16, 17]), which should be explored in future work. It would also be beneficial to the field to determine which coil combination strategies offer the best SNR in newer advanced MRS editing protocols such as Hadamard Encoding and Reconstruction of MEGA-Edited Spectroscopy (HERMES) [18–20] and Hadamard Editing Resolves Chemicals Using Linear-Combination Estimation of Spectra (HERCULES) [21]. Lastly, it will likewise be pertinent to explore this in different voxels of interest that are relevant to clinical populations, such as the prefrontal cortex in major depressive disorder [22], especially since SNR can differ based on the location of the MRS voxel of interest [9].

In conclusion, noise decorrelation methods, especially GLS, produced higher SNR than other methods for both GABA+ and NAA. It also produced lower CVs than most other methods. Although it remains to be determined which noise decorrelation method is best, GLS should be investigated to further optimize SNR in MRS studies, especially since it is relatively easy to implement using existing protocols.

## Data statement

The GABA-edited MEGA-PRESS data are available online in the Big GABA NITRC repository, supported by NIH grant R01EB016089: https://www.nitrc.org/projects/biggaba/

## CRediT statement

Conceptualization: M.M.

Data curation: N/A

Formal analysis: A.E.B. and M.M.

Funding acquisition: M.M.

Investigation: A.E.B. and M.M.

Methodology: N/A

Project administration: N/A

Software: M.M.

Resources: M.M.

Supervision: M.M.

Validation: N/A

Visualization: A.E.B. and M.M.

Writing – original draft: A.E.B. and M.M.

Writing – review and editing: A.E.B. and M.M.

